# SARS-CoV-2 is well adapted for humans. What does this mean for re-emergence?

**DOI:** 10.1101/2020.05.01.073262

**Authors:** Shing Hei Zhan, Benjamin E. Deverman, Yujia Alina Chan

## Abstract

In a side-by-side comparison of evolutionary dynamics between the 2019/2020 SARS-CoV-2 and the 2003 SARS-CoV, we were surprised to find that SARS-CoV-2 resembles SARS-CoV in the late phase of the 2003 epidemic after SARS-CoV had developed several advantageous adaptations for human transmission. Our observations suggest that by the time SARS-CoV-2 was first detected in late 2019, it was already pre-adapted to human transmission to an extent similar to late epidemic SARS-CoV. However, no precursors or branches of evolution stemming from a less human-adapted SARS-CoV-2-like virus have been detected. The sudden appearance of a highly infectious SARS-CoV-2 presents a major cause for concern that should motivate stronger international efforts to identify the source and prevent near future re-emergence. Any existing pools of SARS-CoV-2 progenitors would be particularly dangerous if similarly well adapted for human transmission. To look for clues regarding intermediate hosts, we analyze recent key findings relating to how SARS-CoV-2 could have evolved and adapted for human transmission, and examine the environmental samples from the Wuhan Huanan seafood market. Importantly, the market samples are genetically identical to human SARS-CoV-2 isolates and were therefore most likely from human sources. We conclude by describing and advocating for measured and effective approaches implemented in the 2002-2004 SARS outbreaks to identify lingering population(s) of progenitor virus.

---

Several reports have noted that SARS-CoV-2 appears genetically stable and not under much pressure to adapt, which bodes well for diagnostics, vaccine, and therapeutics development (1–4). How long a particular antiviral, antibody, or vaccine will be effective against SARS-CoV-2 depends greatly on how fast and how extensively the target gene or protein is evolving. To identify effective therapies, immense efforts have already been directed towards elucidating the precise structure of SARS-CoV-2 proteins, ideally, in complex with potential drug candidates - some of which are already undergoing clinical trials (5–11). Any new SARS-CoV-2 variant that can escape or confound these highly precise approaches will subvert efforts to control the pandemic. Therefore, it is very important to ascertain that the SARS-CoV-2 genes targeted in these efforts have stabilized and to identify novel virulent variants as soon as possible.

To gain a better understanding of how stable the SARS-CoV-2 genome is, we first performed a side-by-side comparison of evolutionary dynamics between SARS-CoV-2 and SARS-CoV. For this analysis, we curated high quality genomes spanning ~3-month periods for the following groups: 11 genomes for early-to-mid epidemic SARS-CoV, 32 genomes for late epidemic SARS-CoV, and 46 genomes for SARS-CoV-2 that included an early December, 2019 isolate, Wuhan-Hu-1, and 15 randomly selected genomes from each month of January through March, 2020 sampled from diverse geographical regions (methods in **Supplementary Materials**). We were surprised to find that SARS-CoV-2 exhibits low genetic diversity in contrast to SARS-CoV, which harbored considerable genetic diversity in its early-to-mid epidemic phase (**Figure 1**) (12); nucleotide diversity estimates (13,14) across all sites, non-synonymous and synonymous, for each locus examined are provided in the **Supplementary Table**.

**Figure 1.**
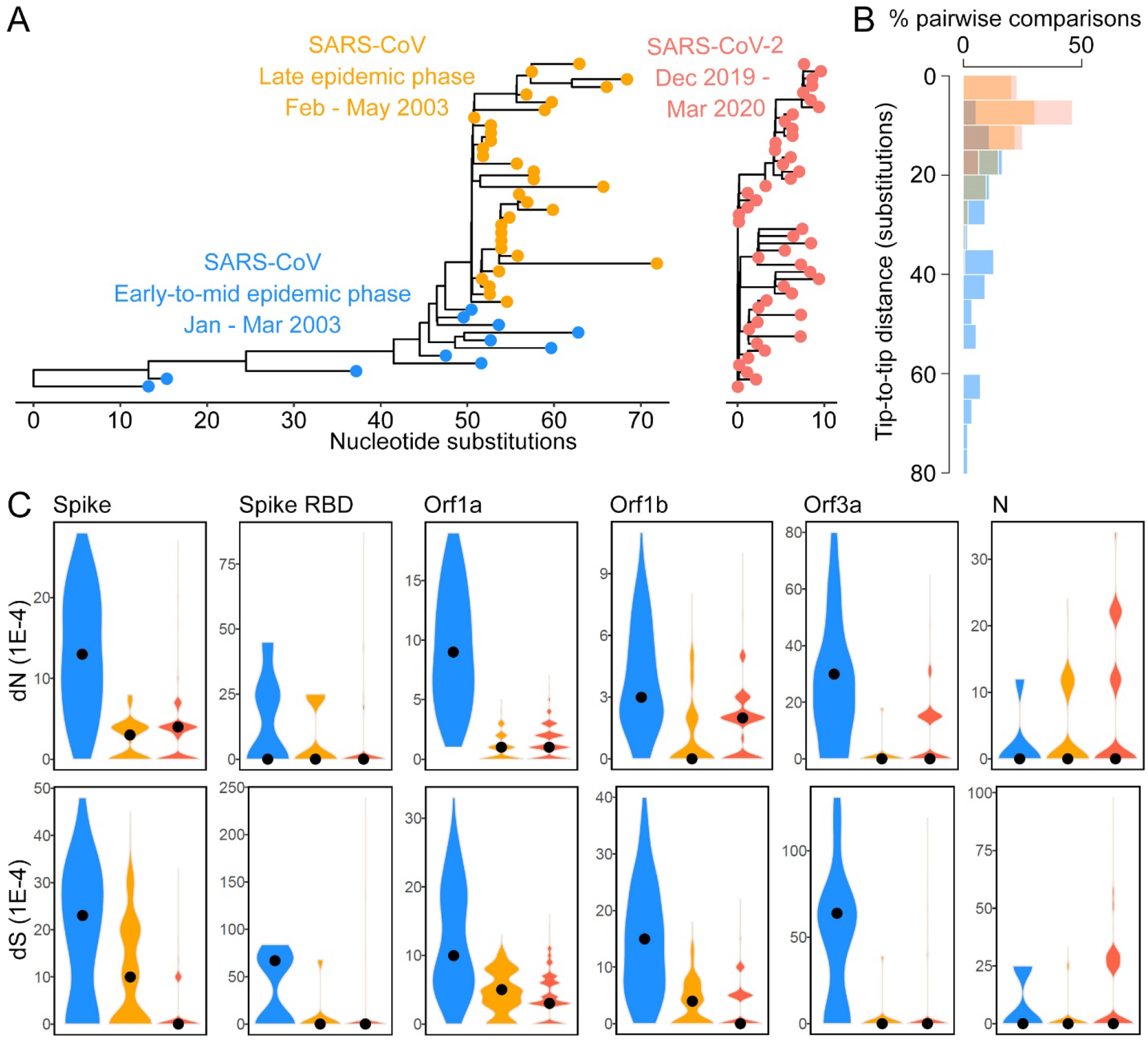
Comparison of the genetic divergence of SARS-CoV and SARS-CoV-2. **(A)** Maximum likelihood trees built with IQtree (19). We curated 11 early-to-mid epidemic SARS-CoV genomes, 32 late epidemic SARS-CoV genomes, and 46 SARS-CoV-2 genomes consisting of a December, 2019 Wuhan-Hu-1 isolate and 15 isolates from each month of January, February, and March, 2020. **(B)** Tip-to-tip distance of each tree: SARS-CoV-2 (red) is less polymorphic than early-to-mid epidemic (blue) SARS-CoV over similar 3-month periods based on the current sampling approach (resampling test, p < 0.01). **(C)** Distribution of pairwise non-synonymous (dN) and synonymous (dS) substitution rates in the Spike, S RBD, Orf1a, Orf1b, Orf3a, and N gene across 151 SARS-CoV-2 genomes: 50 from each month of January, February, and March, 2020, in addition to the Wuhan-Hu-1 isolate.

SARS-CoV was observed to adapt under selective pressure that was highest as it crossed from Himalayan palm civets (intermediate host species) to humans and diminished towards the end of the epidemic (15–18); this series of adaptations between species and in humans culminated in a highly infectious SARS-CoV that dominated the late epidemic phase. In comparison, SARS-CoV-2 exhibits genetic diversity that is more similar to that of late epidemic SARS-CoV (**Figure 1**, **Supplementary Table**). In fact, the exceedingly high level of identity shared among SARS-CoV-2 isolates makes it impractical to model site-wise selection pressure. As more mutations occur and, ideally, when SARS-CoV-2-like viruses from an intermediate host species are identified, it will become possible to model selection pressure as was done for SARS-CoV.

An examination of 43 SARS-CoV and 46 SARS-CoV-2 genomes revealed a striking difference in the number of substitutions over similar 3-month periods (**Figure 1A**), with more genetic polymorphism in early-to-mid epidemic SARS-CoV compared to late epidemic SARS-CoV or SARS-CoV-2 (**Figure 1B**). To rule out a subsampling artefact, we performed one hundred sampling experiments from all high-quality SARS-CoV-2 sequences on GISAID; IQtree phylogenies were built from the one hundred 46-taxon subsampled sequence sets, i.e., 15 randomly selected samples from each month of January through March, 2020 from diverse geographical regions, in addition to Wuhan-Hu-1. The maximum tip-to-tip distance (number of substitutions between two genomes) of the early-to-mid epidemic SARS-CoV tree (~85 substitutions) was greater than that of all one hundred resampled SARS-CoV-2 trees (~15-25 substitutions; p < 0.01). Even by April 28, 2020, the SARS-CoV-2 genomes available on GISAID spanning 4 months exhibited modest genetic diversity (**Supplementary Figure 1**) as compared to early-to-mid epidemic SARS-CoV.

A caveat of this analysis is that genetic diversity can be inflated by sampling from diverse or high traffic locations and be skewed by factors such as effective population size and virus transmission rate; whereas a sampling bias towards isolates sharing a more recent common ancestor will underestimate genetic diversity (20–22). For this reason, we sampled within similar 3-month periods and ensured that there was geographic spread in the sampling. Unfortunately, due to the scarcity of 2003 SARS-CoV samples and information, we cannot ensure that the early-to-mid epidemic samples did not straddle deep splits in the tree. Nonetheless, the SARS-CoV genomes used in our analysis have been used in dozens of studies to examine SARS-CoV adaptive evolution. The division of SARS-CoV cases into early-to-mid versus late phase epidemiologically-linked clusters has been validated (17). Furthermore, the late epidemic SARS-CoV and SARS-CoV-2 samples are from more numerous, international locations compared to early-to-mid epidemic SARS-CoV, which consists of infections only in China. The SARS-CoV outbreak spanning November, 2002 to August, 2003 was estimated to result in 8,422 cases (23) as compared to the more than 850,000 known SARS-CoV-2 cases by the end of March (and more than 3 million cases by the end of April). A considerable portion of SARS-CoV-2 cases are asymptomatic or mild, leading to an underestimation of the virus population size. Yet, early-to-mid epidemic SARS-CoV, which was sampled from a more limited population in a more limited location, exhibits the most genetic diversity.

We proceeded to compare the evolutionary dynamics of SARS-CoV and SARS-CoV-2 in terms of the non-synonymous and synonymous substitution rates (dN and dS) in each gene. Non-synonymous substitutions, as compared to synonymous substitutions, are generally more likely to result in functionally distinct variants. Therefore, dN and dS have commonly been used to model selective pressure on each gene, and were used to determine that the spike (S), Orf3a, and Orf1a genes experienced strong selective pressure in the SARS-CoV epidemic (15–17,24). Importantly, the S protein binds to host receptors and influences host specificity, while the Orf3a-encoded accessory protein facilitates the endocytosis of S (25,26). Orf3a and S have been proposed to share a co-evolutionary relationship (27,28). We sampled 50 SARS-CoV-2 genomes from each month of January, February, and March, 2020 from diverse locations in addition to Wuhan-Hu-1. For the spike (S), Orf3a, and Orf1a genes, the dN and dS in SARS-CoV-2 is more similar to late epidemic than early-to-mid epidemic SARS-CoV (**Figure 1C**, **Figure 2**). In comparison, the highly conserved Orf1b (encodes RNA-dependent RNA polymerase RdRp and helicase Hel), which did not undergo strong positive selection in SARS-CoV (15), exhibits similarly low dN across the three CoV groups (**Figure 1C**, **Figure 2**).

**Figure 2.**
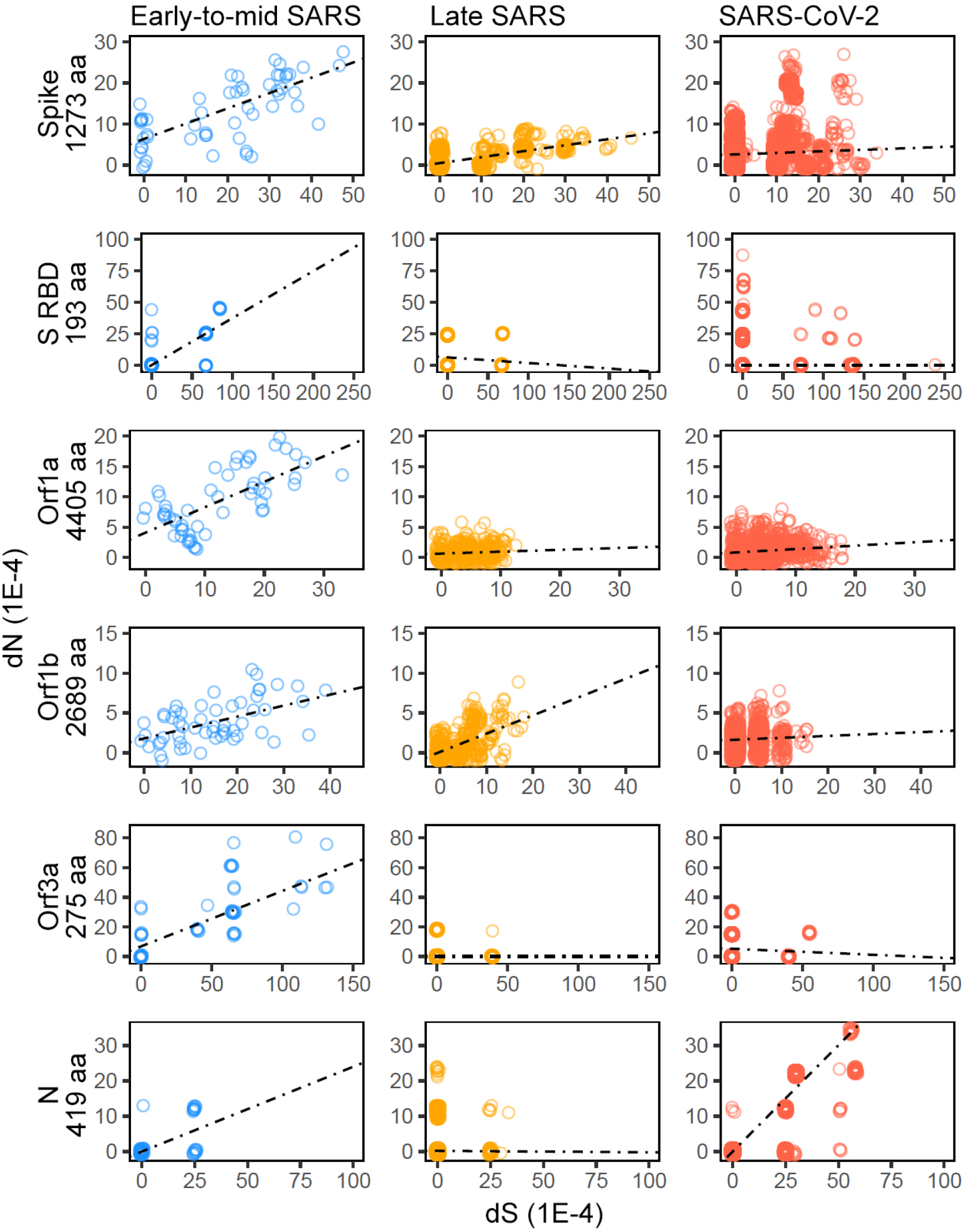
Pairwise dN and dS in each indicated gene in early-to-mid epidemic SARS-CoV, late epidemic SARS-CoV, and SARS-CoV-2. Sizes of each encoded protein in SARS-CoV-2 are provided. Importantly, the number of points per plot is influenced by the number of genomes analyzed (11 early-to-mid, 32 late epidemic SARS-CoV, 151 SARS-CoV-2) and cannot be used to infer the quantity of substitutions; for that, please refer to **Figures 1**, **3**, and **4**. The slopes estimated under robust regression using M-estimation (35) are shown by the dashed lines.

In consideration that several therapies and antibodies in development target the SARS-CoV-2 S, it is important to track non-synonymous substitutions and predict the evolution of resistance. We analyzed the non-synonymous substitutions that occurred in the S of SARS-CoV and SARS-CoV-2 over the course of each epidemic. Numerous adaptive mutations that evolved in SARS-CoV S RBD have been experimentally demonstrated to enhance binding to the human ACE2 receptor and facilitate cross-species transmission, e.g., residues N479 and T487 (29,30), as well as K390, R426, D429, T431, I455, N473, F483, Q492, Y494, R495 (31); or predicted to have been positively selected, e.g. residues 239, 244, 311, 479, 778 (17) (**Figure 3**). In contrast, the majority of the non-synonymous substitutions in SARS-CoV-2 S are distributed across the gene at low frequency and have not been reported to confer adaptive benefit (**Figure 4**). Yet, the SARS-CoV-2 S has been demonstrated to bind more strongly to human ACE2 and has a superior plasma membrane fusion capacity compared to the SARS-CoV S (32,33). The only site of notable entropy in the SARS-CoV-2 S, D614G, lies outside of the RBD and is not predicted to impact the structure or function of the protein (34). Its prevalence in international COVID-19 cases has been attributed to the substitution occurring early in the pandemic leading to a founder’s effect. There is no evidence of a more virulent strain of SARS-CoV-2 emerging despite passage through more than 3 million human hosts by the time of this analysis.

**Figure 3.**
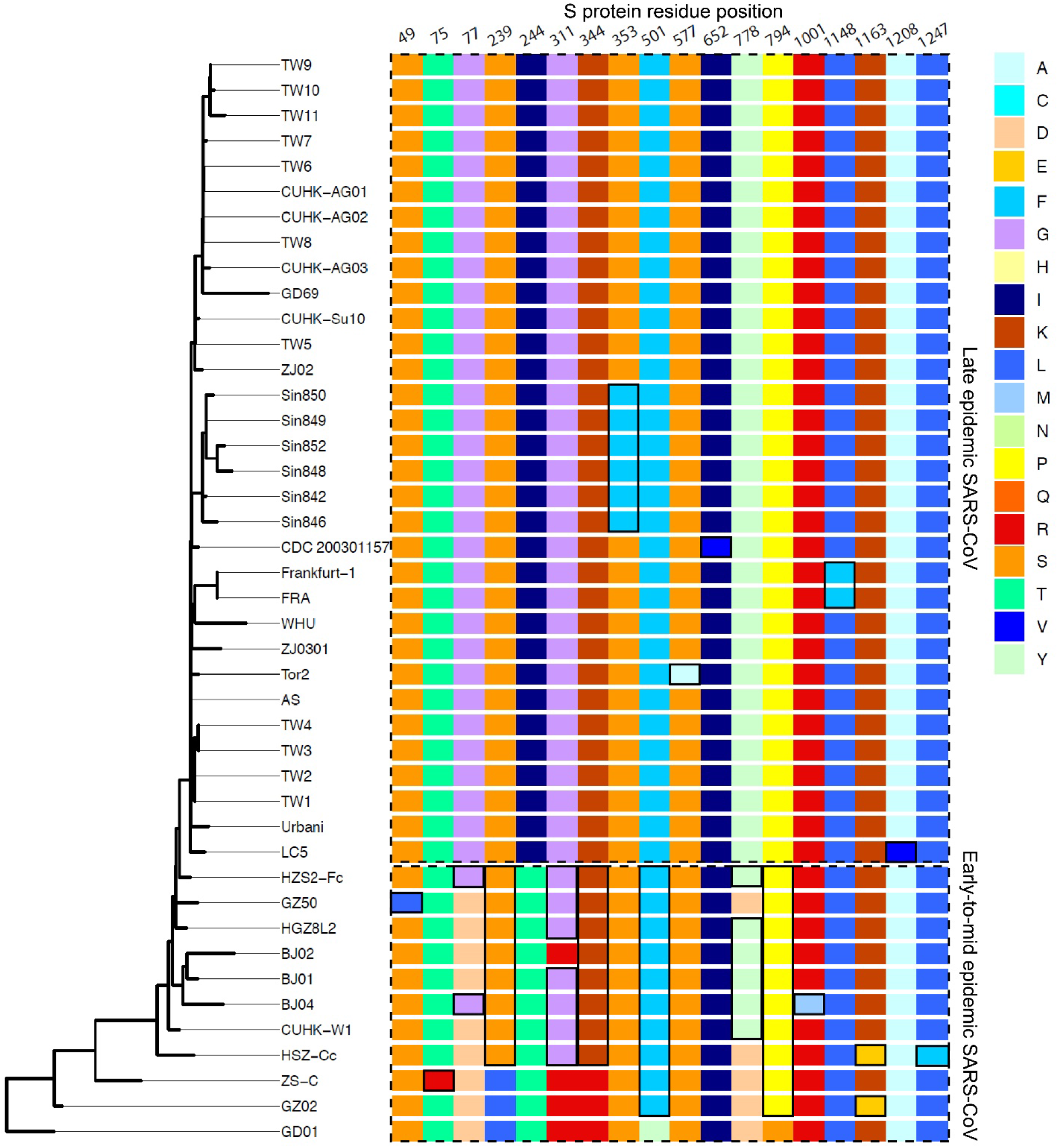
Spike protein evolution in SARS-CoV. Only residues that vary among the isolates are shown. The more recent variants at each position in each epidemic phase are emphasized with black outlines. Residues 75, 239, 244, 311, 778, 1148, and 1163 have been predicted to be positively selected during the interspecies epidemic (17); civet variants not shown.

**Figure 4.**
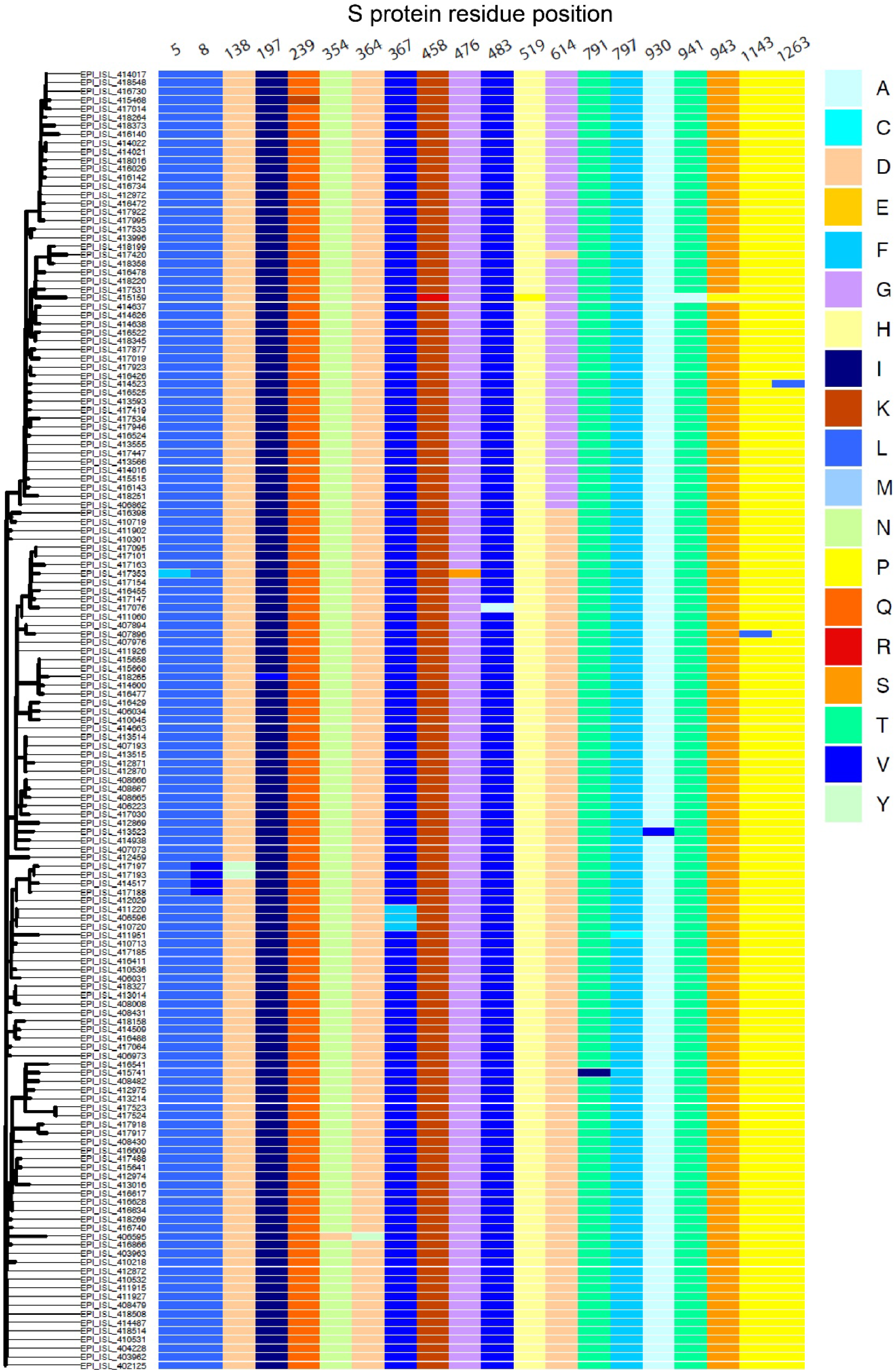
Spike protein evolution in SARS-CoV-2. The 151 isolates in analysis include Wuhan-Hu-1 (EPI_ISL_402125, bottom row) and 50 randomly selected genomes per month between January through March, 2020. Only residues that vary among the isolates are shown.

The pairwise comparisons of dN and dS, alongside a dearth of signs of emerging adaptive mutations, suggest that by the time SARS-CoV-2 was first detected in late 2019, it was already well adapted for human transmission to an extent more similar to late epidemic than to early-to-mid epidemic SARS-CoV. One possible scenario is that the SARS-CoV-2 outbreak in late 2019 resulted from a bottleneck event similar to the late epidemic SARS-CoV cases that stemmed from a single superspreader who visited Metropole Hotel in Hong Kong, China in late February, 2003 (36). In comparison to the SARS-CoV epidemic, the SARS-CoV-2 epidemic appears to be missing an early phase during which the virus would be expected to accumulate adaptive mutations for human transmission. However, if this were the origin story of SARS-CoV-2, there is a surprising absence of precursors or branches emerging from a less recent, less adapted common ancestor among humans and animals. In the case of SARS-CoV, the less human-adapted SARS-CoV progenated multiple branches of evolution in both humans and animals (**Figure 1**, **Figure 5**). In contrast, SARS-CoV-2 appeared without peer in late 2019, suggesting that there was a single introduction of the human-adapted form of the virus into the human population. This has important implications regarding the risk of SARS-CoV-2 re-emergence in the near future and the severity of its consequences.

**Figure 5.**
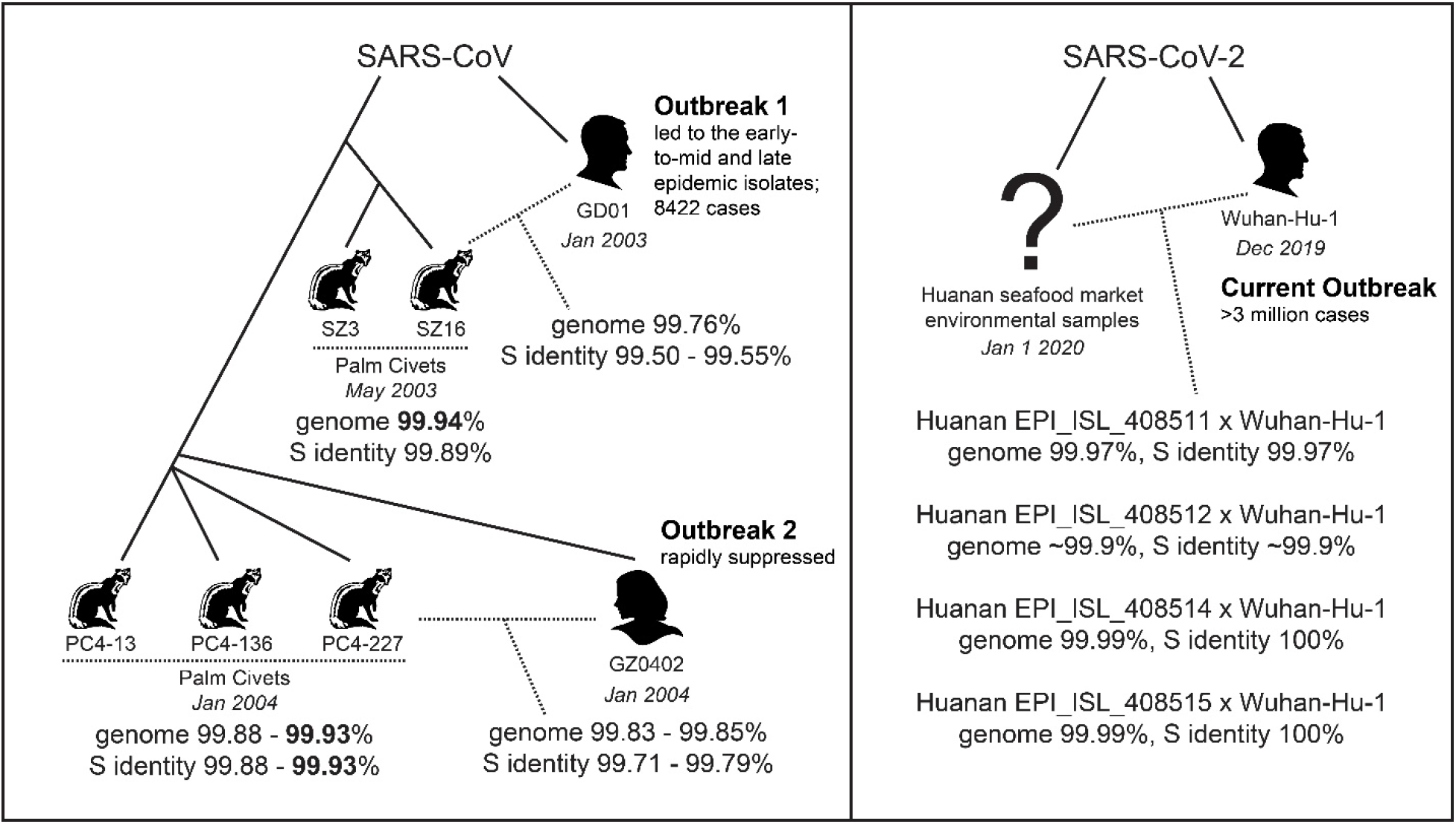
Comparison of genome and S gene identity across human, animal, and environmental samples. Dotted lines indicate the samples under comparison for which genome or S gene identity are shown. Due to ambiguous bases and lack of data regarding Huanan seafood market environmental sample quality, it is difficult to assess shared identity with high accuracy. This was particularly true for sample *EPI_ISL_408512*, which had frequent stretches of undetermined bases in its genome sequence. Shared identity above 99.9% is bolded in the left panel depicting SARS-CoV isolates. Only isolates from within the same species in a narrow window of time shared 99.9% S gene identity, i.e., only civets sampled at the same time shared >99.9% S gene identity; the civet isolates from May, 2003 (SZ3 and SZ16) and January, 2004 (PC4-13, PC4-136, and PC4-227) shared only up to 99.71% genome and 99.42% gene S identity.

It is important to recall that there were two SARS-CoV outbreaks in 2002-2004, each arising from separate palm civet-to-human transmission events (**Figure 5**): the first emerged in late 2002 and ended in August, 2003; the second arose in late 2003 from a lingering population of SARS-CoV progenitors in civets. The second outbreak was swiftly suppressed due to diligent human and animal host tracking, informed by lessons from the first outbreak (37,38). To prevent similar consecutive outbreaks of SARS-CoV-2 today, it is vital to learn from the past and implement measures to minimize the risk of additional SARS-CoV-2-like precursors adapting to and re-emerging among humans. To do so, it is important to identify the route by which SARS-CoV-2 adapted for human transmission. However, there is presently little evidence to definitively support any particular scenario of SARS-CoV-2 adaptation. Did SARS-CoV-2 transmit across species into humans and circulate undetected for months prior to late 2019 while accumulating adaptive mutations? Or was SARS-CoV-2 already well adapted for humans while in bats or an intermediate species? More importantly, does this pool of human-adapted progenitor viruses still exist in animal populations? Even the possibility that a non-genetically-engineered precursor could have adapted to humans while being studied in a laboratory should be considered, regardless of how likely or unlikely (39).

## What is known about possible intermediate hosts and SARS-CoV-2 species tropism?

Speculations that pangolins are the likely intermediate animal host stemmed from the discovery of a pangolin CoV that shares 95.4% S amino acid identity and six key RBD residues with SARS-CoV-2 (40). Since then, another closely related lineage of pangolin CoVs has been identified (41). However, the unique polybasic furin cleavage site in the SARS-CoV-2 S is not found in pangolin CoVs (42), and SARS-CoV-2 is not a recent recombinant involving any of the CoVs sampled to date (41,43,44). The CoV that is most closely related to SARS-CoV-2 is RaTG13, a bat CoV that was identified at the Wuhan Institute of Virology and originally isolated from the Yunnan Province of China (45). RaTG13 shares 96.2% genome identity with the Wuhan-Hu-1 SARS-CoV-2 isolate. In comparison, the most closely related pangolin CoV MP789 shares only 84.1% and 84.0% genome identity with Wuhan-Hu-1 and RaTG13, respectively. No evidence as yet points to the adaptation of SARS-CoV-2 for human infection in pangolins or the transmission of SARS-CoV-2 from pangolins to humans.

In addition, it is plausible for SARS-CoV-2 S to have evolved its broad species tropism naturally in bats or a wide range of intermediate species. The SARS-CoV-2 S is predicted to bind to ACE2 from potentially more than 100 diverse species (46–48), and was demonstrated to bind more strongly than the SARS-CoV S to ACE2 from both bat and human (33). The S of RaTG13 is also capable of binding to human ACE2 although the virus does not infect humans (49). Similarly, the S of human MERS-CoV was found to bind to receptors from humans, camels, and bats, and could adapt to semi-permissive host receptors within three passages in cell culture (50). Therefore, although no sampled bat CoVs have been found to possess a SARS-CoV-2-like S RBD, these findings collectively suggest that some CoVs in nature are evolving S that can bind at an optimal level to the same receptor across diverse species (43), potentially by interfacing with highly conserved parts of the receptor. As other groups have recommended, CoV sampling from more species - to avoid bias stemming from the focused scrutiny of Malayan pangolins - will provide us with a better grasp of the range of species that harbor CoVs with similar RBDs to SARS-CoV-2, as well as the natural diversity of bat CoVs (43).

There has been considerable debate among scientists and the public on whether SARS-CoV-2 originated from the Wuhan Huanan seafood market (2). According to the Chinese CDC’s website, accessed on April 27, 2020, SARS-CoV-2 was detected in environmental samples at the Huanan seafood market, and the Chinese CDC suggested that the virus originated from animals sold there (51). However, phylogenetic tracking suggests that SARS-CoV-2 had been imported into the market by humans (52). To look for clues regarding an intermediate animal host, we turned to samples collected from the market in January, 2020. In contrast to the thorough and swift animal sampling executed in response to the 2002-2004 SARS-CoV outbreaks to identify intermediate hosts (37,53), no animal sampling prior to the shut down and sanitization of the market was reported. Details about the sampling are sparse: 515 out of 585 samples are environmental samples, and the other 70 were collected from wild animal vendors; it is unclear whether the latter samples are from animals, humans, and/or the environment. Only 4 of the samples, which were all environmental samples from the market, have passable coverage of SARS-CoV-2 genomes for analysis. Even so, these contain ambiguous bases that confound genetic clustering with human SARS-CoV-2 genomes. Nonetheless, the market samples did not form a separate cluster from the human SARS-CoV-2 genomes. We compared the market samples to the human Wuhan-Hu-1 isolate, and discovered >99.9% genome identity, even at the S gene that has exhibited evidence of evolution in previous CoV zoonoses. In the SARS-CoV outbreaks, >99.9% genome or S identity was only observed among isolates collected within a narrow window of time from within the same species (**Figure 5**) (15). The human and civet isolates of the 2003/2004 outbreak, which were collected most closely in time and at the site of cross-species transmission, shared only up to 99.79% S identity (**Figure 5**) (37). It is therefore unlikely for the January market isolates, which all share 99.9-100% genome and S identity with a December human SARS-CoV-2, to have originated from an intermediate animal host, particularly if the most recent common ancestor jumped into humans as early as October, 2019 (54,55). The SARS-CoV-2 genomes in the market samples were most likely from humans infected with SARS-CoV-2 who were vendors or visitors at the market. If intermediate animal hosts were present at the market, no evidence remains in the genetic samples available.

## Conclusion

The lack of definitive evidence to verify or rule out adaptation in an intermediate host species, humans, or a laboratory, means that we need to take precautions against each scenario to prevent re-emergence. We would like to advocate for measured and effective approaches to identify any lingering population(s) of SARS-CoV-2 progenitor virus, particularly if these are similarly adept at human transmission. The response to the first SARS-CoV outbreak deployed the following strategies that were key to detecting SARS-CoV adaptation to humans and cross-species transmission, and could be re-applied in today’s outbreak to swiftly eliminate progenitor pools: (i) Sampling animals from markets, farms, and wild populations for SARS-CoV-2-like viruses (38). (ii) Checking human samples banked months before late 2019 for SARS-CoV-2-like viruses or SARS-CoV-2-reactive antibodies to detect precursors circulating in humans (56). In addition, sequencing more SARS-CoV-2 isolates from Wuhan, particularly early isolates if they still exist, could identify branches originating from a less human-adapted progenitor as was seen in the 2003 SARS-CoV outbreak. It would be curious if no precursors or branches of SARS-CoV-2 evolution are discovered in humans or animals. (iii) Evaluating the over-or underrepresentation of food handlers and animal traders among the index cases to determine if SARS-CoV-2 precursors may have been circulating in the animal trading community (57). While these investigations are conducted, it would be safer to more extensively limit human activity that leads to frequent or prolonged contact with wild animals and their habitats.

## Supporting information

GISAID Acknowledgments

## Acknowledgments

We thank contributors of SARS-CoV-2 genomes (GISAID) and SARS-CoV genomes (ViPR); Fusion Genomics Corporation and Compute Canada for providing clusters; Dr. Mohammad Qadir (CEO/CSO), Greg Stazyk (CTO) of Fusion Genomics Corporation, and Dr. Matthew Solomonson (Broad Institute) for reviewing this work; and Dr. Shao-Lun Liu (Associate Professor, Tunghai University) for PAML advice. We thank SHZ’s supervisor, Dr. Sarah P. Otto, Canada Research Chair in Theoretical and Experimental Evolution, the University of British Columbia for her generous mentorship in supporting and reviewing this work.

## Supplementary Materials and Methods

### Data curation and random sampling of SARS-CoV-2 genome sequences

We analyzed genome sequences at least 29,000 bases in length of SARS-CoV from the 2002-2003 outbreak (early-to-mid phase and late phase) and SARS-CoV-2 from the 2019-2020 pandemic. The sequences of SARS-CoV and SARS-CoV-2 were downloaded from ViPR (1) and GISAID (2), respectively. Contributors of the SARS-CoV-2 genome sequences are listed in our GISAID Acknowledgments file. The sequences from cruise ship patients were excluded. The sequences from the earliest part of the COVID-19 pandemic (December, 2019 in Wuhan) were excluded because they were of poor quality (3). Additionally, SARS-CoV-2 sequences were removed from the selection pool if they contained undetermined or ambiguous bases after the ends were trimmed. The SARS-CoV-2 sequences were randomly selected such that the same number of sequences were taken from each month of collection and, within each month, the sequences were evenly sampled from each geographic region; the probability of each particular sequence being sampled was 1/(number of sequences in country X * total number of countries). Using this sampling strategy, we derived sequence sets that are temporally and spatially well represented.

### Phylogenetic tree reconstruction

Multiple sequence alignments were built for the SARS-CoV sequences and SARS-CoV-2 sequences separately using MAFFT version 7.453 (4) with the ‘--auto’ option. Then, phylogenetic trees were reconstructed using IQtree version 1.6.12 (5) with automatic model selection using ModelFinder (6). The GD01 (GenBank: AY278489) and Wuhan-Hu-1 (GISAID: EPI_ISL_402125) sequences were used to root the SARS-CoV and SARS-CoV-2 genome phylogenies, respectively. The GD01 sequence was obtained from one of the early patients of the 2003 outbreak and carries the 29-nt Orf8 region found in palm civet hosts of SARS-CoV but was lost during human-to-human transmission of the virus (7). The Wuhan-Hu-1 sequence is a high-quality genome sequence from an early patient of the COVID-19 pandemic (8). The phylogenetic trees were processed and visualized using the R packages *ape* version 5.3 (9), *adephylo* version 1.1-11 (10), and *ggtree* version 2.0.4 (11).

### Estimation of pairwise dN and dS

Nonsynonymous and synonymous substitution rate (dN and dS) were estimated under pairwise maximum-likelihood comparisons (12) using codeml, which is part of the PAML version 4.8 package (13). The SARS-CoV sequence set was split into two sets corresponding to the early-to-mid phase and the late phase of the 2003 outbreak. Indels were not considered in our analysis (see Supplementary Discussion on *Indels and Orf8 deletion*). It is currently impractical to assess the site-wise selective pressure on SARS-CoV-2 genes due to the low genetic diversity. In particular, smaller genes are less amenable to dN and dS analysis within short time frames with limited genetic diversity. Nonetheless, we have plotted additional dN and dS plots in **Supplementary Figures 2**-**4**, but these should be interpreted with caution. Mainly, adding the most closely related animal CoVs to the analysis did not change our findings and potentially added a skew in the case of bat CoV RaTG13, which shares only ~96.2% genome identity with SARS-CoV-2 isolates (**Supplementary Figures 2**-**4**).

In **Supplementary Figures 2 and 4**, the following closely related bat and civet CoVs were included in the analysis: **For SARS-CoV, civet CoVs:** SZ16 (AY304488), SZ3 (AY304486), HC/GZ/32/03 (AY545918), HC/GZ/81/03 (AY545917), HC/SZ/266/03 (AY545916), HC/SZ/79/03 (AY545914), and HC/SZ/DM1/03 (AY545915). **For SARS-CoV-2, a bat CoV:** RaTG13 (MN996532.1).

## Supplementary Text

### Analysis of environmental samples from the Huanan seafood market in Wuhan

We found available sequence data for five environmental samples collected on January 1, 2020, from the Huanan seafood market. Unfortunately, the quality of these samples cannot be ascertained because the raw sequencing data is not available. Single nucleotide differences or differences near the ends of fragments could stem from low sequencing quality or coverage. We compared the environmental sample sequences to the human Wuhan-Hu-1 SARS-CoV-2 genome reference from December, 2019 (8):

Sample *EPI_ISL_408511*, with sequences totaling 28,557-nt, shared 99.97% genome identity and 99.97% S gene identity with Wuhan-Hu-1. Sample *EPI_ISL_408512*, with sequences totaling 25,342-nt, shared 99.87% genome identity with Wuhan-Hu-1. The fragments covering 3585-nt of the 3822-nt S gene were ~99.9% identical. Sample *EPI_ISL_408513* had a 439-nt sequence from its S gene, which was 100% identical to the Wuhan genome reference. The other ~600-nt in fragments were visibly low in quality and mapped to Orf1b, a highly conserved region in SARS-like-CoVs. It was not useful to analyze these low-quality short sequences. Samples *EPI_ISL_408514 and 408515* each had sequences totaling 29,891-nt sharing 99.99% genome identity and 100% S gene identity with Wuhan-Hu-1. The two samples were 99.99% identical to each other.

Many of the single nucleotide differences came from ambiguous base calls, for example, R or Y, in the environmental sample sequences. If these were excluded from our alignments, the percent identity between the environmental samples and Wuhan-Hu-1 would increase. However, the lack of access to the raw sequencing data prevents us from assessing the reliability of the sequence similarities or disparities. There is also no information about what these environmental samples were derived from, e.g., type of shop (live-animal, meat), species of animals in that area, type of surface the sample was collected from, or whether COVID-19 human cases frequented that area before the samples were collected on January 1, 2020.

### Indels and Orf8 deletion

The SARS-CoV 29-nt deletion or the deletion of Orf8 was found to be detrimental to virus replication across mammalian cell lines, including primate and human cell lines (14). Muth et al. postulated that “the 29 nt deletion in SARS-CoV is the result of a founder effect that has permitted survival in spite of reduction of fitness”. In agreement with these findings, the Chinese SARS Molecular Epidemiology Consortium found that “Major deletions were observed in the Orf8 region of the genome, both at the start and the end of the epidemic” (15). In their report, Muth et al. also discussed the relationship between reduced virulence or replication, prolonged survival of infected animals, and consequences for viral propagation during an epidemic (14): a less replication-efficient (possibly asymptomatic) virus may have an advantage when quarantining measures are in place by hiding and re-emerging after social distancing measures have been lifted. The impact of deletions in Orf8 is especially important in consideration of the recent discovery of a 382-nt deletion in Orf8 of SARS-CoV-2 (16). However, in consideration of the complexity associated with deletions in Orf8 of SARS-CoVs, we opted to exclude the analysis of indels in our work due to the inability to discern whether these indels could be perpetuated due to founder effects rather than a true adaptive benefit.

---

**Supplementary Table.** Nucleotide diversity estimates among isolates for each ORF.

|  | **SARS-CoV**<br>**early-to-mid** | **SARS-CoV**<br>**late** | **SARS-CoV-2**<br>**(n=46)** | **SARS-CoV-2**<br>**(n=151)** |
| --- | --- | --- | --- | --- |
| Spike | 5.3091 | 1.6230 | 0.7478 | 0.9992 |
| Spike RBD | 1.1273 | 0.4355 | 0.0435 | 0.1452 |
| Orf1a | 13.0909 | 3.6215 | 2.5546 | 2.5920 |
| Orf1b | 5.2000 | 1.3908 | 1.4290 | 1.5873 |
| Orf3a | 2.5818 | 0.1250 | 0.6531 | 0.5840 |
| N | 0.5091 | 0.3710 | 1.1275 | 1.1539 |
| **Non-synonymous sites** |  |  |  |  |
| Spike | 3.8545 | 0.6230 | 0.6174 | 0.7885 |
| Spike RBD | 0.6182 | 0.3145 | 0.0000 | 0.1187 |
| Orf1a | 9.2000 | 1.8581 | 1.2087 | 1.3528 |
| Orf1b | 2.2909 | 0.5954 | 0.9981 | 0.9254 |
| Orf3a | 1.7818 | 0.0625 | 0.6531 | 0.5310 |
| N | 0.1818 | 0.3085 | 0.8783 | 0.8698 |
| **Synonymous sites** |  |  |  |  |
| Spike | 1.4545 | 1.0000 | 0.1304 | 0.2107 |
| Spike RBD | 0.5091 | 0.1210 | 0.0435 | 0.0265 |
| Orf1a | 3.8909 | 1.7634 | 1.3459 | 1.2392 |
| Orf1b | 2.9091 | 0.7954 | 0.4309 | 0.6619 |
| Orf3a | 0.8000 | 0.0625 | 0.0000 | 0.0530 |
| N | 0.3273 | 0.0625 | 0.2493 | 0.2842 |

**Supplementary Figure 1.**
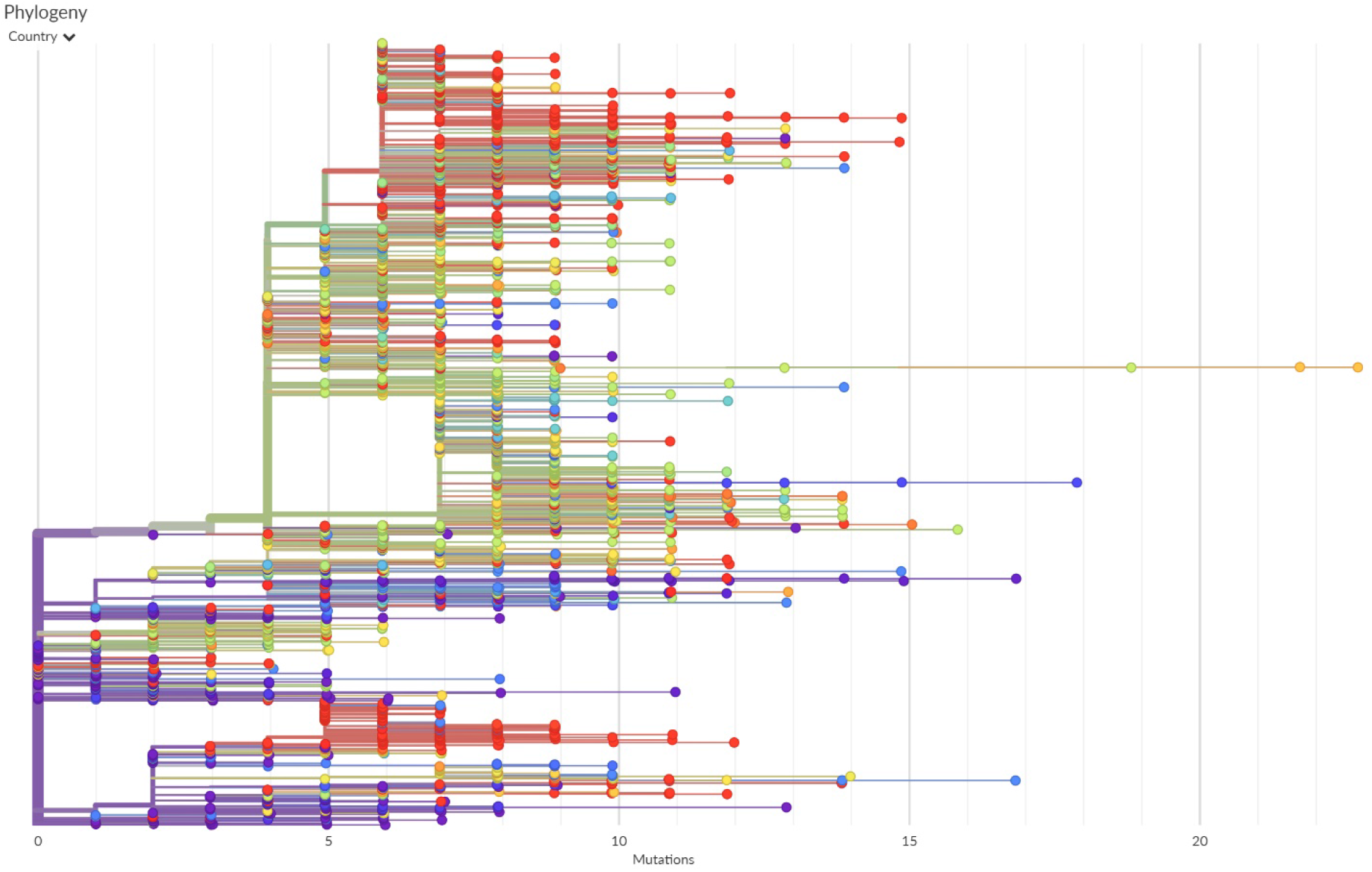
Divergence of SARS-CoV-2 genomes rooted with the Wuhan-Hu-1 genome reference. This figure was obtained from NextStrain (17) on April 28, 2020, and describes the genetic divergence spanning ~4 months between the Wuhan-Hu-1 reference collected in December, 2019, and the genomes collected up to late April.

**Supplementary Figure 2.**
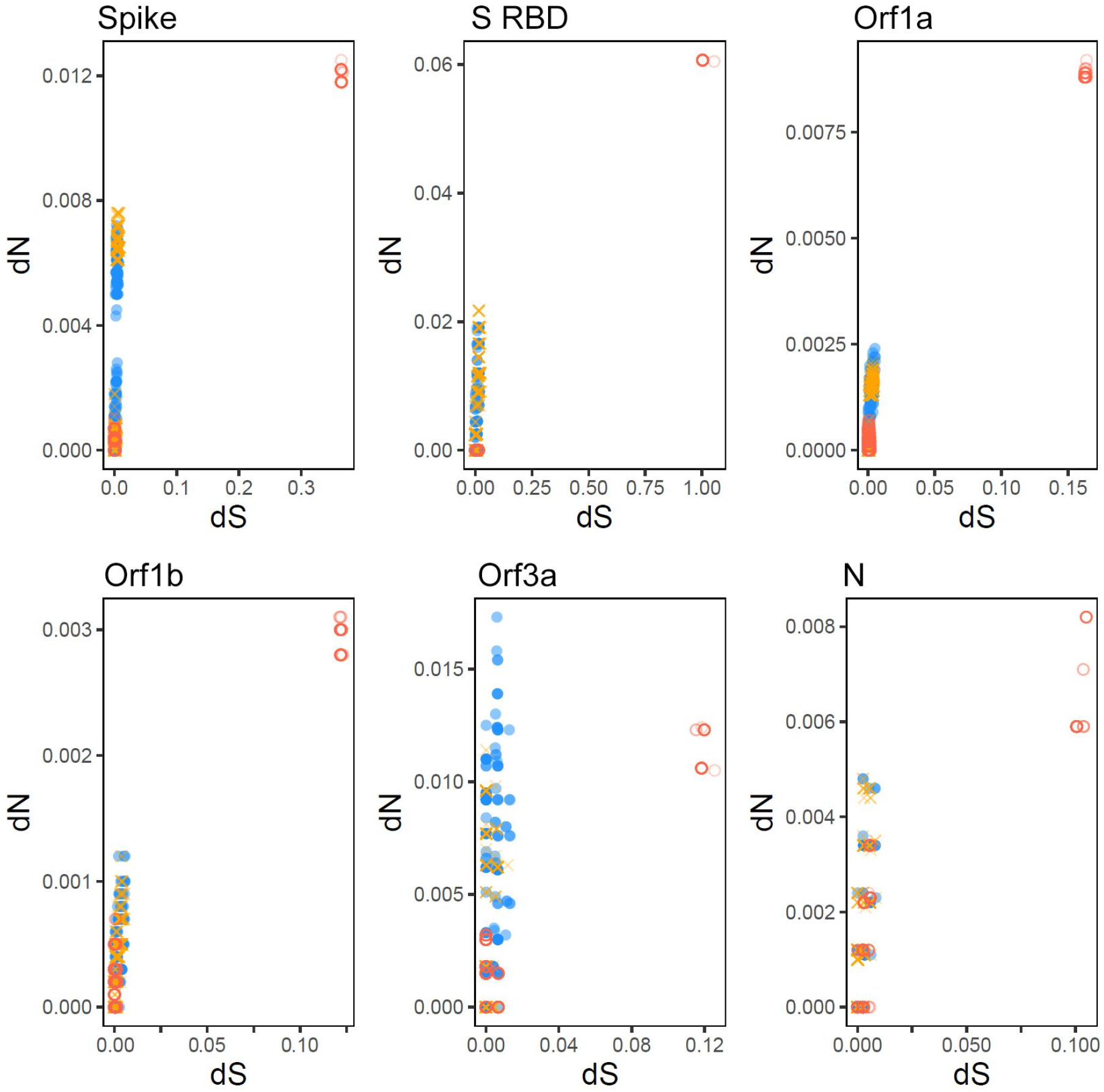
Plots of pairwise dN versus dS for each locus examined. SARS-CoV early-to-mid epidemic (11 genomes), late epidemic (32 genomes), and SARS-CoV-2 (46 genomes) are represented in blue (colored circle), orange (cross), and red (empty circle), respectively. The bat CoV RaTG13 was used in the calculation of dN and dS for SARS-CoV-2; seven 2003 civet SARS-CoVs were used in the calculation of dN and dS for early-to-mid and late epidemic SARS-CoV. We did not include pangolin CoV MP789 in the analysis for SARS-CoV-2 due to (i) low 84.1% genome identity and only 86.4% nucleotide identity at the S RBD despite 96.4% amino acid identity, and (ii) stretches of ambiguous sequence that prohibited locus-specific dN and dS estimations.

**Supplementary Figure 3.**
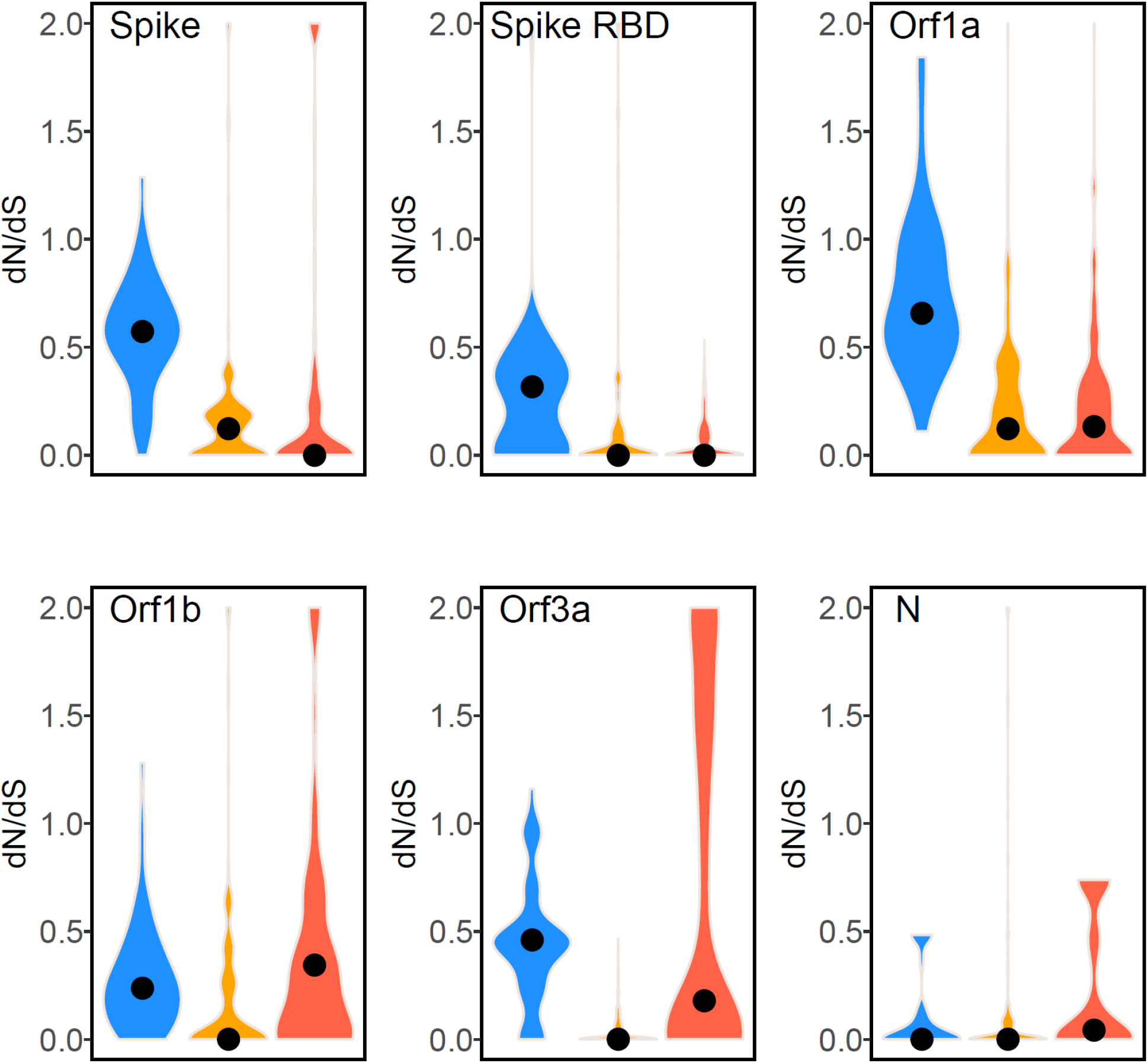
Pairwise ratios of non-synonymous to synonymous substitution rate (dN/dS) in the Spike, S RBD, Orf1a, Orf1b, Orf3a, and N. High estimates (>2) were removed as per common practice. The analysis for SARS-CoV-2 is based on 46 genomes: Wuhan-Hu-1 and 15 randomly selected genomes from each month of January, February, and March, 2020. No intermediate animal host CoVs have been identified for SARS-CoV-2. Thus, single population dN/dS are shown as opposed to inter-species dN/dS in order to keep the values for SARS-CoV and SARS-CoV-2 comparable. The same plots using bat CoV RaTG13 and civet SARS-CoVs are shown in **Supplementary Figure 4**.

**Supplementary Figure 4.**
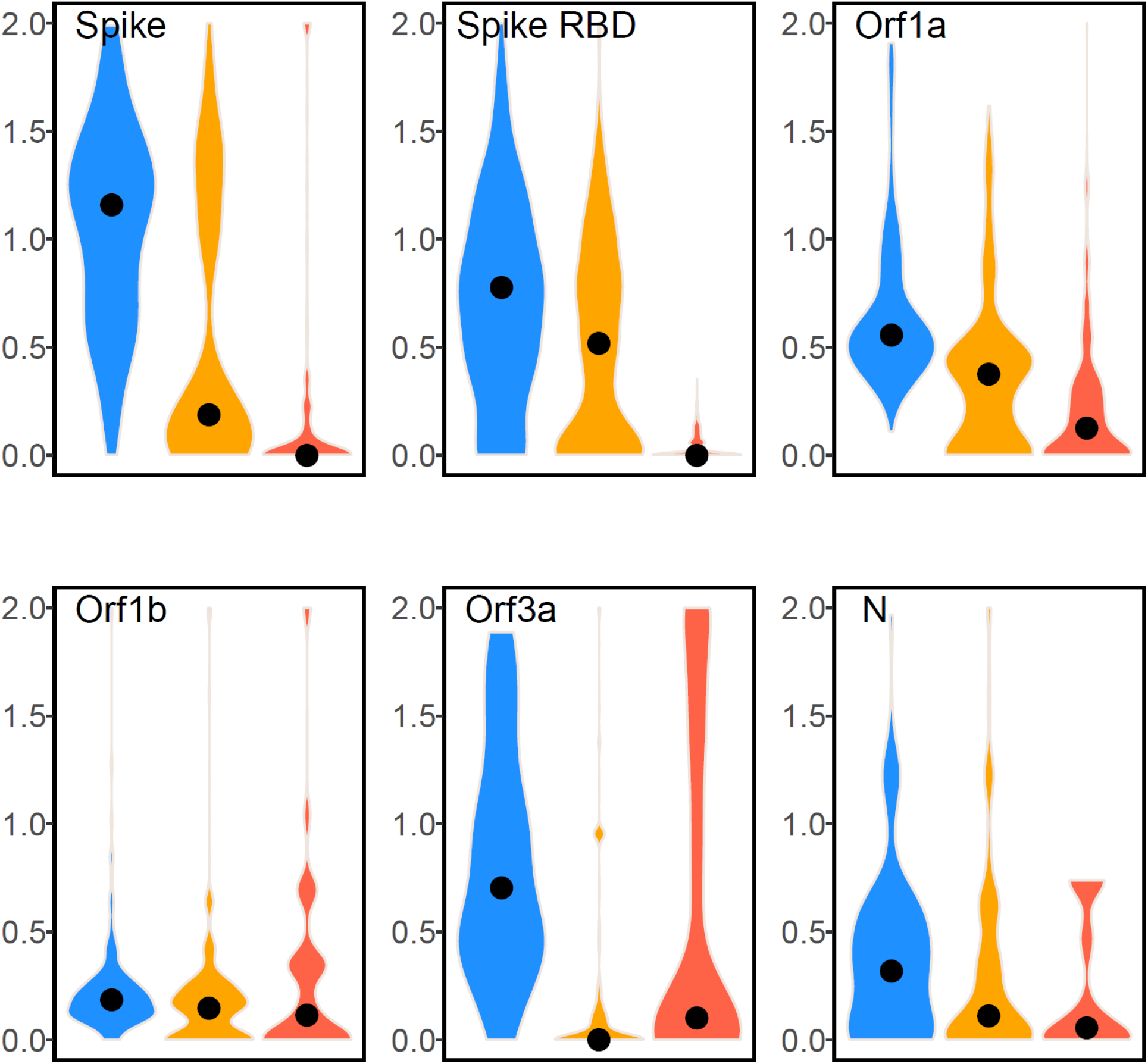
Pairwise ratios of non-synonymous to synonymous substitution rate (dN/dS) in the Spike, S RBD, Orf1a, Orf1b, Orf3a, and N. High estimates (>2) were removed as per common practice. The analysis for SARS-CoV-2 is based on 46 genomes: Wuhan-Hu-1 and 15 randomly selected genomes from each month of January, February, and March, 2020. No intermediate animal host CoVs have been identified for SARS-CoV-2. The dN/dS calculations were performed using the most closely related animal CoVs: palm civet SARS-CoVs from 2003 (99.76% genome identity) alongside the human SARS-CoVs, and bat CoV RaTG13 (96.2% genome identity) alongside the human SARS-CoV-2s.

